# Using genotype data to distinguish pleiotropy from heterogeneity: deciphering coheritability in autoimmune and neuropsychiatric diseases

**DOI:** 10.1101/030783

**Authors:** Buhm Han, Jennie G Pouget, Kamil Slowikowski, Eli Stahl, Cue Hyunkyu Lee, Dorothee Diogo, Xinli Hu, Yu Rang Park, Eunji Kim, Peter K Gregersen, Solbritt Rantapää Dahlqvist, Jane Worthington, Javier Martin, Steve Eyre, Lars Klareskog, Tom Huizinga, Wei-Min Chen, Suna Onengut-Gumuscu, Stephen S Rich, Major Depressive Disorder Working Group of the Psychiatric Genomics Consortium, Naomi Wray, Soumya Raychaudhuri

## Abstract

Shared genetic architecture between phenotypes may be driven by a common genetic basis (pleiotropy) or a subset of genetically similar individuals (heterogeneity). We developed BUHMBOX, a well-powered statistical method to distinguish pleiotropy from heterogeneity using genotype data. We observed a shared genetic basis between 11 of 17 tested autoimmune diseases and type I diabetes (T1D, p<10^−12^) and 11 of 17 tested autoimmune diseases and rheumatoid arthritis (RA, p<10^−7^). This sharing could not be explained by heterogeneity (corrected p_BUHMBOX_>0.2 using 6,670 T1D cases and 7,279 RA cases), suggesting that shared genetic features in autoimmunity are due to pleiotropy. We observed a shared genetic basis between seronegative and seropostive RA (p<10^−22^), explained by heterogeneity (p_BUHMBOX_=0.008 in 2,406 seronegative RA cases). Consistent with previous observations, we observed genetic sharing between major depressive disorder (MDD) and schizophrenia (p<10^−9^). This sharing is not explained by heterogeneity (p_BUHMBOX_=0.28 in 9,238 MDD cases).

## INTRODUCTION

Recent studies have demonstrated that many diseases share risk alleles^1-4^ and exhibit significant coheritability^5-7^. Traditional approaches for detecting coheritability include twin or family studies^8, 9^. Now alternative approaches using genome-wide association study (GWAS) data from unrelated individuals have been developed. Polygenic risk score approaches^3, 10, 11^ build genetic risk scores (GRSs) for one phenotype and test their association with a second phenotype. Mixed-model approaches^5, 6, 12^ can estimate the genetic covariance between two traits on the observed scale. From the genetic covariance one can also calculate the genetic coheritability and genetic correlation^6^. Cross-trait LD Score regression utilizes linkage disequilibrium (LD) and summary statistics obtained from GWAS to estimate genetic correlation attributable to SNPs^7^. In addition, the p-values of independent SNPs associated with multiple phenotypes can be tested for a significant deviation from the null distribution^2^. These approaches have been applied to demonstrate significant shared genetic structure among many phenotypes^5, 7, 13^ including autoimmune^2^ and neuropsychiatric diseases^3, 6, 11^. Coheritability and genetic sharing suggests the possibility of *pleiotropy,* defined here as the sharing of risk alleles across traits at specific genetic variants or at a genome-wide level. Pleiotropy can occur when the same variant causes different diseases (*biological pleiotropy,* e.g. variant R620W in *PTPN22* is associated with multiple autoimmune diseases)^14^, or when a variant causes development of a phenotype that then drives the development of a second phenotype (*mediated pleiotropy,* e.g. rare coding region variants in *LDLR* that increase LDL cholesterol levels are associated with increased risk of myocardial infarction)^15^.

However, it remains uncertain whether the observed shared genetic structure is the consequence of true pleiotropy, or the consequence of heterogeneity. Here, we define *heterogeneity* as the situation where a patient cohort consists of genetically distinct subgroups that may or may not result in distinct symptom profiles and treatment outcomes. This type of heterogeneity can occur in the context of misclassifications (e.g. cases with atypical presentation for a different disease are erroneously included), molecular subtypes (e.g. a subset of cases share pathogenesis with a different disease, possibly as a result of biological or mediated pleiotropy), or ascertainment bias (e.g. cases also affected with a different disease are more likely to come to clinical attention and be included in the study). These situations can result in a subgroup of cases that is genetically similar to another disease, creating shared genetic structure. Indeed, there is mounting evidence that misclassifications^16-19^, etiological diversity^20^, and ascertainment bias^21^ are prevalent across certain human diseases, leading to the conclusion that significant heterogeneity may exist^22-25^. Since the potential contribution of heterogeneity to any genetic sharing observed between diseases represents a critical component of predictive medicine, there is a need for statistical methods to detect heterogeneity on the basis of commonly available genetic data.

## RESULTS

### Overview of BUHMBOX

Genetic sharing between two diseases, disease A (D_A_) and disease B (D_B_), could be due to pleiotropy, but could also be due to heterogeneity (i.e. some D_A_ cases are genetically more similar to D_B_ cases). If we calculated GRSs for D_A_ cases using D_B_-associated loci and their effect sizes (GRS_B_), GRS_B_ would be associated with D_A_ case status under either pleiotropy or heterogeneity. Under pleiotropy, some D_B_ risk alleles impose D_A_ risk, and D_B_ risk alleles will be enriched in D_A_ cases compared to controls. Under heterogeneity, a subset of D_A_ cases will have genetic characteristics of D_B_, and therefore D_B_ risk alleles will also be enriched in those individuals. In both situations, the enriched D_B_ risk alleles in D_A_ cases will result in significant associations between D_A_ status and GRS_B_.

To detect heterogeneity, even in the presence of pleiotropy, we developed BUHMBOX (Breaking Up Heterogeneous Mixture Based On Cross-locus correlations). BUHMBOX leverages the fact that in the setting of heterogeneity, D_B_ risk alleles are enriched only in a specific subset of D_A_ cases while in true pleiotropy, D_B_ risk alleles are enriched uniformly across the entire set of D_A_ cases (**Figure 1**). BUHMBOX tests for enrichment differences of D_B_ risk alleles among D_A_ cases by estimating correlations between independent loci. If D_B_ risk alleles are enriched in one subgroup, the expected correlations of risk allele dosages between loci will be consistently positive (for details see **Supplementary Table 1** and **Supplementary Information**).

**Figure 1.**
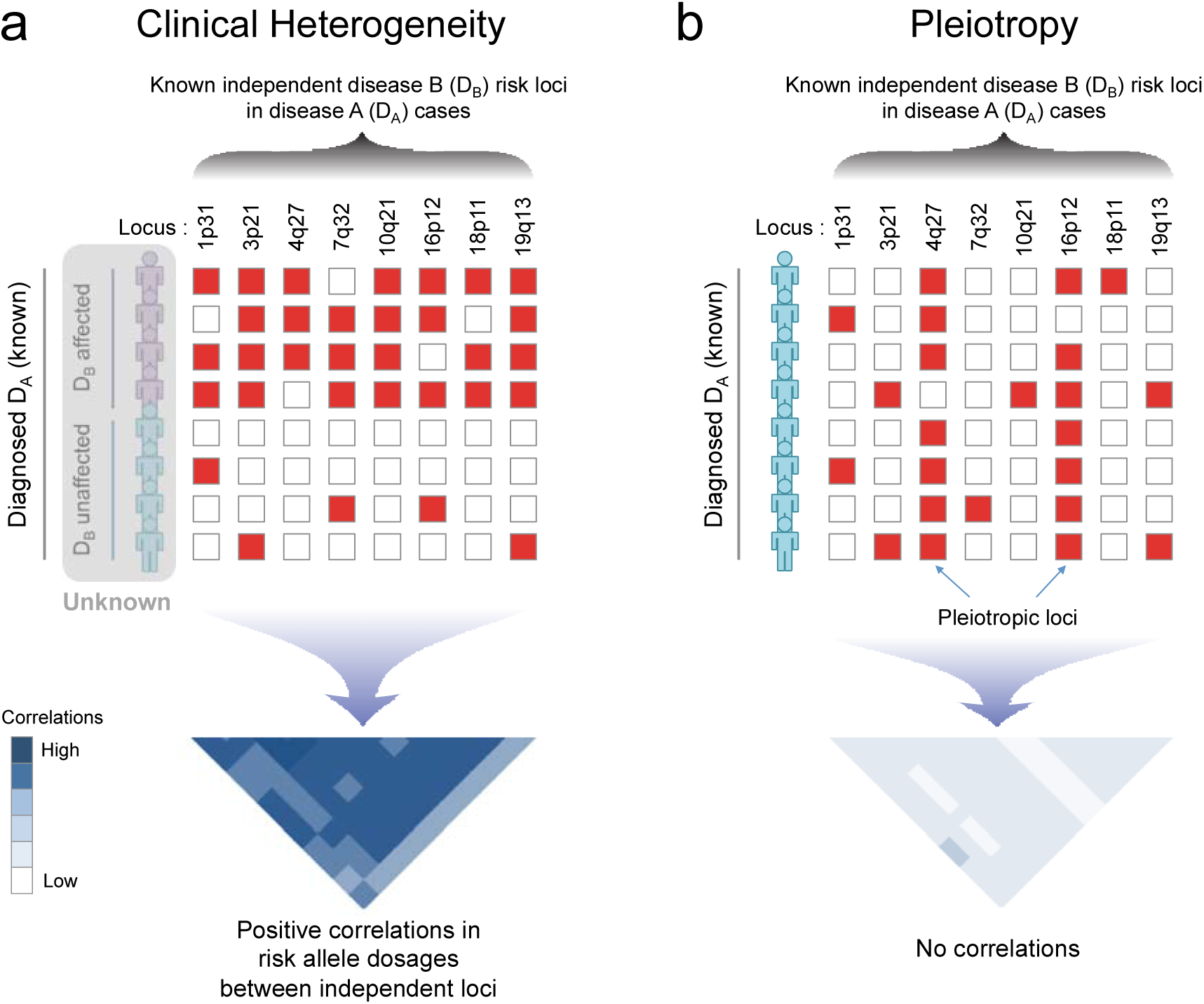
Overview of BUHMBOX. (a) Under the scenario of heterogeneity, risk alleles of disease B (D_B_)-associated loci will be enriched in a subgroup of disease A (D_A_) cases, producing positive correlations between Db risk allele dosages from independent loci. (b) Under the scenario where there is no heterogeneity, but D_A_ and D_B_ share alleles due to pleiotropy, D_B_ risk alleles will be uniformly distributed, and have no correlations. Red boxes: risk alleles; white boxes: non-risk alleles.

### BUHMBOX discriminates between heterogeneity and pleiotropy

We wanted to demonstrate that BUHMBOX detects heterogeneity, but is robust to the presence of pleiotropy. To this end, we conducted simulations with the following parameters: sample size of D_A_ case individuals (*N*), number of risk loci associated to D_B_ (*M*), and the proportion of D_A_ cases that actually show genetic characteristics of D_B_ (heterogeneity proportion, or *π*). To simulate realistic distributions of effect sizes and allele frequencies, we sampled odds ratio (OR) and risk allele frequency (RAF) pairs from reported associations in the GWAS catalog^28^ (**Methods**).

To characterize the false positive rate (FPR) of BUHMBOX we simulated 1,000,000 studies (*N*=2,000 and *M*=50) where there was neither heterogeneity (*π*=0, **Methods**) or pleiotropy. BUHMBOX obtained appropriate false positive rates at all statistical significance thresholds evaluated (p<0.05 to 0.0005, **Supplementary Table 2**); for example, at p<0.05 we observed a 5.1% FPR.

To evaluate the FPR of BUHMBOX where there actually was pleiotropy without heterogeneity (*π*=0), we simulated 1,000 studies (*N*=2,000 and *M*=50) where we assumed D_A_ and D_B_ shared 10% of risk loci. We quantified the proportion of instances where BUHMBOX and GRS approaches obtained p-values smaller than the threshold p<0.05. With the presence of true pleiotropy without heterogeneity, GRS appropriately demonstrated 87.6% power to detect shared genetic structure. BUHMBOX demonstrated an appropriate false positive rate of 4.6% (**Supplementary Figure 1**).

Finally, to evaluate BUHMBOX’s power to detect heterogeneity we repeated the above simulations assuming there was no pleiotropy, but that there was indeed subtle heterogeneity. Specifically we assumed that 10% of D_A_ cases were actually D_B_ (*π*=0.1). Here, BUHMBOX demonstrated 91.1% power to detect heterogeneity at p<0.05 (**Supplementary Figure 1**). The GRS approach demonstrated 100% power to detect shared genetic structure.

Together, these simulations illustrate that BUHMBOX is sensitive to heterogeneity but robust to the presence of pleiotropy, while the GRS detects both scenarios and cannot discriminate between the two. BUHMBOX complements existing methods for detecting pleiotropy by helping to interpret shared genetic structure identified with these approaches (**Supplementary Table 1**).

### Weighting SNPs by their effect sizes increases power

BUHMBOX combines multiple correlations into one statistic. In order to maximize power, we defined a scheme to weight the correlations between alleles as a function of their effect sizes and allele frequencies (**Methods**). In simulations we observed substantial power gain with this weighting scheme. Assuming 1,000 cases and 50 loci, we compared the power of BUHMBOX implemented with and without weighting correlations (equation (12) in **Supplementary Information**). Across a wide range of *π* values we observed that the weighting scheme in BUHMBOX dramatically increased power (**Figure 2**). For example, at *π*=0.1 the weighted implementation of BUHMBOX obtained 74% compared to the unweighted implementation which obtained only 36% power.

**Figure 2.**
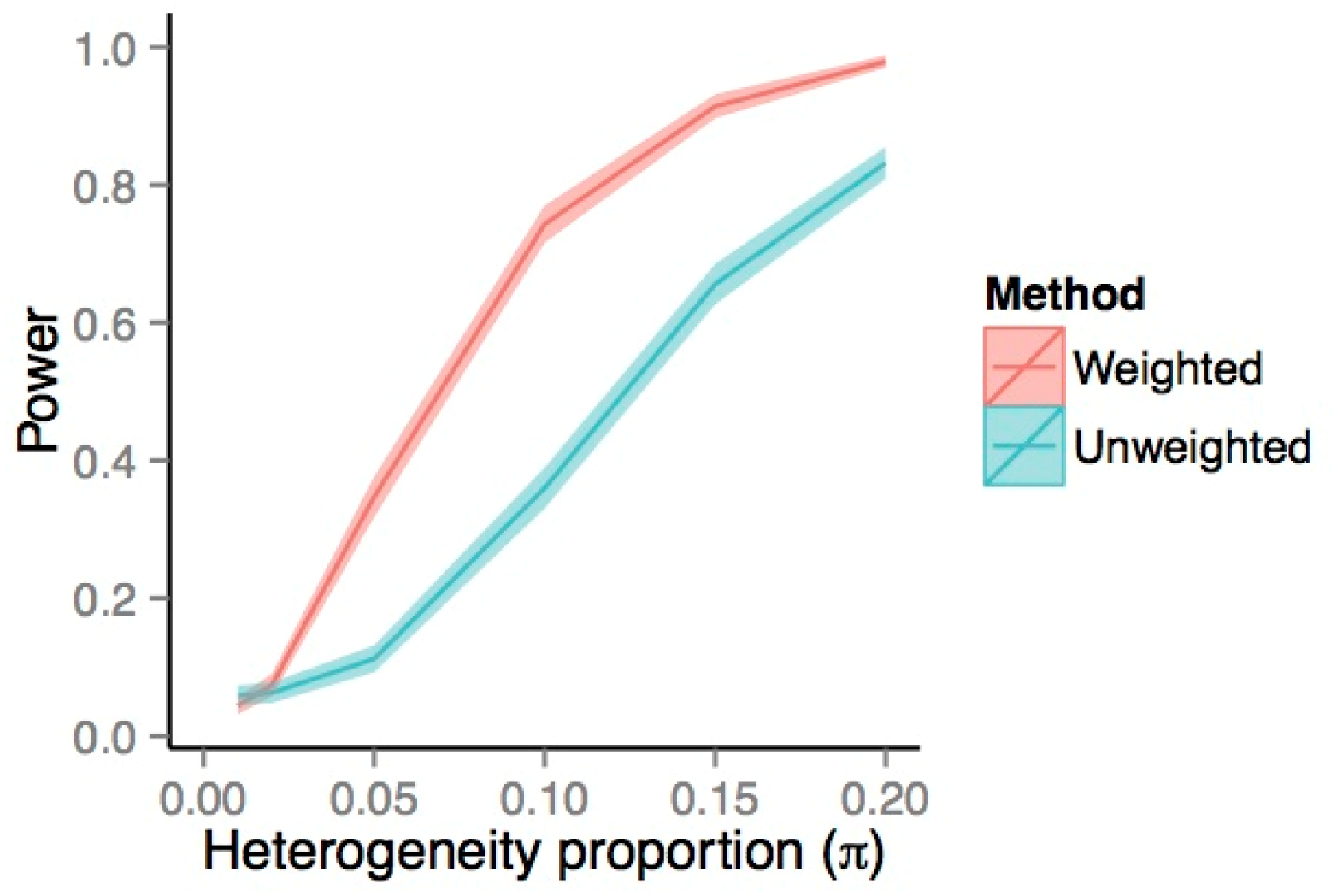
Power gain by weighting SNPs by allele frequency and effect size. We compared power of BUHMBOX with a weighting scheme that optimally weights correlations between SNPs (weighted) to an alternative approach that weights correlations uniformly (unweighted; equation (12) in **Supplementary Information**). We simulated 1,000 case individuals and assumed 50 risk loci, whose OR and RAFs were sampled from GWAS catalog. The colored bands denote 95% confidence intervals of power estimates. The weighting scheme of BUHMBOX offers a clear power advantage.

### Statistical power as a function of numbers of samples and loci

We benchmarked the statistical power of BUHMBOX under a range of different conditions. Power is a function of many factors including sample size *N* of the cases we are testing for heterogeneity in, number of loci *M* for the coheritabile disease, heterogeneity proportion *π,* RAF, and OR. We sampled pairs of RAF and OR from the GWAS catalog. Given a sample size of *N*=2,000 cases and 2,000 controls, assuming *π*=0.2, our method achieved 92% power at p<0.05 level if we had 50 risk loci (**Figure 3**). As many GWAS now collect more than 2,000 cases, and an increasing number of diseases are approaching 50 known associated loci^26^, BUHMBOX is currently well powered to detect a moderate amount heterogeneity (*π*=0.2) for many human traits. Modest heterogeneity is more challenging to detect at this sample size; power decreased to 67% at *π*=0.1 and to 38% at *π*=0.05. Power can be augmented with larger sample size (**Figure 3**). Power can also be increased by including large numbers of loci with nominal evidence of association in the coheritable disease in addition to established genome-wide significant loci (**Supplementary Figure 2**).

**Figure 3.**
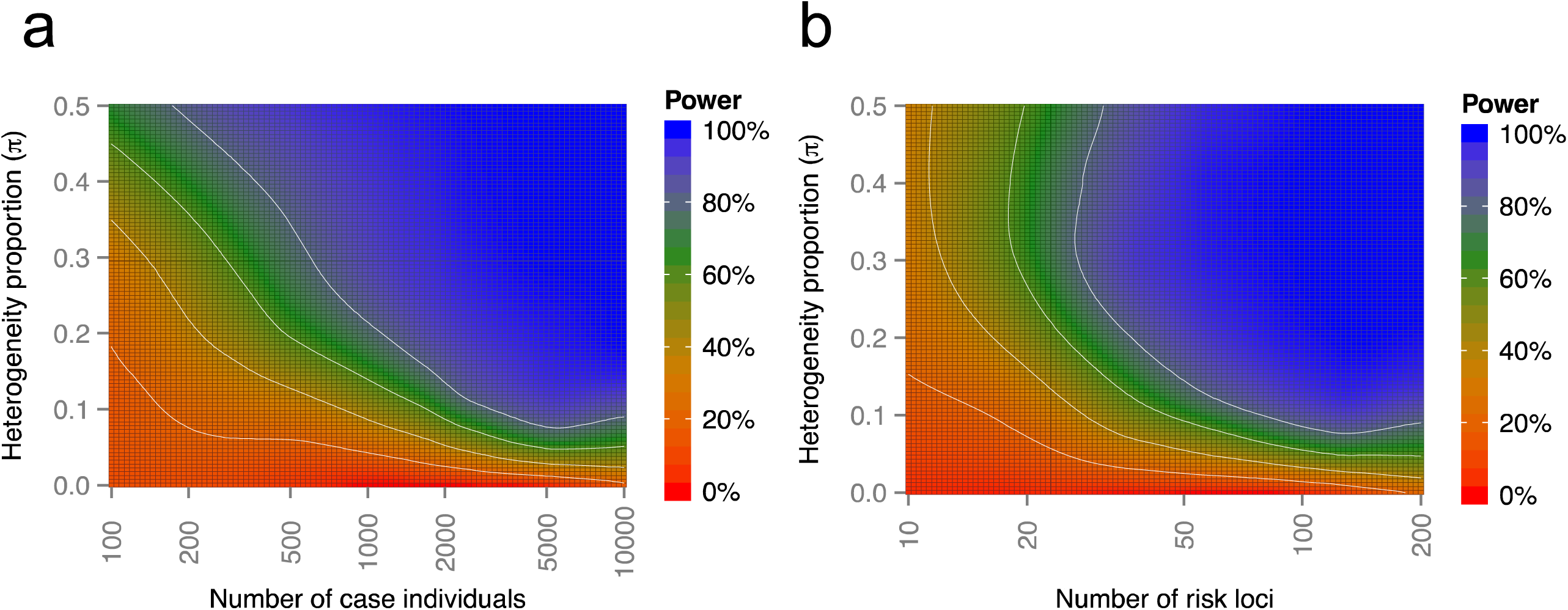

### Controlling for linkage disequilibrium

Although BUHMBOX adequately controlled the FPR when loci were truly independent, we were concerned that long-range LD between two apparently independent loci may introduce false positives^27^. To ensure that BUHMBOX was robust to the effects of LD, we implemented the following strategies in BUHMBOX: (1) stringent LD-pruning of the set of D_B_ loci to exclude SNPs within 1Mb of each other and those with r^2^>0.1, and (2) accounting for any residual LD after pruning by assessing the relative increase of correlations in cases compared to controls (*delta-correlations*). We evaluated the effectiveness of these strategies by measuring FPR using the RA Immunochip Consortium data^28^. We generated 1,000 different loosely pruned (*r*^2^<0.5) SNP sets using the Sweden EIRA data (**Methods**) and measured the FPR without using delta-correlations. As expected, we observed a high FPR (25.2%) at p<0.05. However, when we repeated simulations using stringent pruning (*r*^2^<0.1) and delta-correlations, we were able to conservatively control the FPR (FPR=0.022) at p<0.05.

### Accounting for population stratification

Another potential confounding factor that can challenge independence across loci is population stratification. If population stratification exists, weak correlations between unlinked loci may occur, leading to inappropriate significance. If similar population stratification exists in cases and controls, the use of delta-correlations mitigates this effect. Additionally, to more aggressively control for the effect of stratification at the individual level, we implemented BUHMBOX to regress out PCs from risk allele dosages before calculating correlation statistics. To evaluate the effectiveness of this strategy, we simulated a dataset with extreme population stratification using HapMap^29^ data (60 CEU and 60 YRI founders as cases, and 90 JPG+CHB founders as controls; *λ*_GC_=26.5). As expected, when we randomly sampled 5,000 sets of independent SNPs we observed an inflated BUHMBOX FPR (14.1% at p<0.05). After regressing the effect of ten PCs from risk allele dosages, we observed that the FPR was appropriately controlled (5.7% at p<0.05). As an additional test with more realistic levels of stratification, we merged genotype data from Northern Europe (Sweden EIRA cohort; 2,762 cases/1,940 controls) and Southern Europe (Spain cohort; 807 cases/399 controls) in the RA Immunochip Consortium case-control dataset^28^ (**Methods**) to create a highly stratified dataset. We then randomly sampled 1,000 sets of independent SNPs from this sample. We observed an inflation of the FPR (8.6% at p<0.05), which was appropriately corrected (5.9% at p<0.05) when we regressed out the effect of ten PCs.

### Application to autoimmune diseases

Autoimmune diseases share risk SNPs at many genetic loci^2, 4, 30-34^, clustering in specific immune pathways^2, 25, 34^. We used the GRS approach to evaluate the extent of genetic sharing between autoimmune diseases at a genome-wide level, and then applied BUHMBOX to assess if any observed genetic overlap was due to either true pleiotropy or heterogeneity. We obtained individual-level genotype data from the Type 1 Diabetes Genetics Consortium (T1DGC) UK case-control cohort (6,670 cases and 9,416 controls)^35^ and the RA Immunochip Consortium’s six RA case-control cohorts (7,279 seropositive RA cases and 15,870 controls)^28^ (for sample details, see **Methods**). With these data, we evaluated the genetic sharing between a spectrum of autoimmune diseases with T1D and RA. We obtained associated independent loci for all 18 autoimmune diseases (r^2^<0.1, including MHC SNPs) from ImmunoBase (http://www.immunobase.org, **Supplementary Table 3**), and tested the association of GRSs for these autoimmune diseases with T1D and RA case status.

Unsurprisingly, we observed substantial genetic sharing between autoimmune diseases. In particular T1D demonstrated significant overlap with alopecia areata (AA), autoimmune thyroid disease (ATD), celiac disease (CEL), Crohn’s disease (CRO), juvenile idiopathic arthritis (JIA), primary biliary cirrhosis (PBC), primary sclerosing cholangitis (PSC), RA, Sjogren’s syndrome (SJO), systemic lupus erythematosus (SLE), and vitiligo (VIT) (positive association, p<10^−12^). RA exhibited significant overlap with AA, ankylosing spondylitis (AS), ATD, CEL, JIA, PBC, PSC, SLE, systemic sclerosis (SSC), T1D and VIT (p<10^−7^). Overall, GRSs showed significant positive associations for 11 autoimmune diseases each in T1D and RA cohorts, respectively (GRS p<2.9×10^−3^ [=0.05/17 to correct for 17 diseases tested]; **Table 1, Supplementary Table 4**). We considered only these traits for subsequent analyses.

**Table 1.** Summary of genetic overlap using GRS and BUHMBOX. Only the traits that have significant GRS P-values in positive directions are shown. Significant GRS P-value indicates evidence of shared genetic structure; significant BUHMBOX P-value indicates evidence of heterogeneity. See **Supplementary Table 4** for the full results for all traits tested.

| Cohort data | Test trait | #SNP | GRS P-value | GRS Beta<br>(estimates $\pi$<br>assuming<br>no pleiotropy) | BUHMBOX<br>P-value | BUHMBOX<br>power at<br>$\pi=20\%$ |
| --- | --- | --- | --- | --- | --- | --- |
| T1D | AA | 10 | 1.0E-276 | 0.76 (0.69 – 0.82) | 0.83 | 0.15 |
|  | ATD | 7 | 9.8E-72 | 0.48 (0.40 – 0.56) | 0.30 | 0.05 |
|  | CEL | 38 | 1.6E-80 | 0.32 (0.27 – 0.38) | 0.16 | 0.50 |
|  | CRO | 119 | 2.4E-11 | 0.08 (0.04 – 0.11) | 0.54 | 0.99 |
|  | JIA | 22 | 2.6E-316 | 0.44 (0.40 – 0.47) | 0.37 | 0.96 |
|  | PBC | 19 | 2.9E-28 | 0.16 (0.11 – 0.20) | 0.18 | 0.82 |
|  | PSC | 12 | 3.6E-59 | 0.38 (0.31 – 0.45) | 0.91 | 0.08 |
|  | RA | 68 | 9.1E-204 | 0.55 (0.49 – 0.60) | 0.45 | 0.40 |
|  | SJO | 7 | 2.4E-319 | 0.53 (0.49 – 0.57) | 0.84 | 0.66 |
|  | SLE | 16 | 1.0E-191 | 0.44 (0.39 – 0.48) | 0.79 | 0.91 |
|  | VIT | 12 | 5.1E-207 | 0.59 (0.53 – 0.65) | 0.14 | 0.33 |
| RA | AA | 10 | 5.7E-51 | 0.28 (0.22 – 0.34) | 0.71 | 0.23 |
|  | AS | 24 | 3.9E-08 | 0.10 (0.04 – 0.15) | 0.19 | 0.20 |
|  | ATD | 7 | 2.1E-45 | 0.34 (0.27 – 0.41) | 0.57 | 0.08 |
|  | CEL | 38 | 6.5E-45 | 0.21 (0.17 – 0.26) | 0.57 | 0.63 |
|  | JIA | 22 | 2.3E-286 | 0.36 (0.33 – 0.39) | 0.61 | 0.99 |
|  | PBC | 19 | 3.1E-30 | 0.15 (0.11 – 0.19) | 0.83 | 0.90 |
|  | PSC | 12 | 3.9E-31 | 0.24 (0.18 – 0.31) | 0.46 | 0.12 |
|  | SLE | 16 | 4.3E-13 | 0.10 (0.05 – 0.14) | 0.34 | 0.96 |
|  | SSC | 5 | 1.7E-21 | 0.22 (0.15 – 0.29) | 0.08 | 0.09 |
|  | T1D | 53 | 2.2E-441 | 0.43 (0.40 – 0.46) | 0.29 | 1.00 |
|  | VIT | 12 | 1.8E-25 | 0.18 (0.12 – 0.23) | 0.02 | 0.41 |
| Seronegative RA | Seropositive RA | 14 | 1.1E-23 | 0.30 (0.21 – 0.39) | 0.008 | 0.26 |
| MDD | SCZ | 90 | 1.5E-10 | 0.19 (0.13 – 0.25) | 0.28 | 0.60 |

AA, Alopecia areata; AS, Ankylosing spondylitis; ATD, Autoimmune thyroid disease; CEL, celiac disease; CRO, Crohn's disease; JIA, juvenile idiopathic arthritis; MS, multiple sclerosis; PBC, primary biliary cirrhosis; PSC, primary sclerosing cholangitis; SJO, Sjogren's syndrome; SLE, systemic lupus
erythematosus; SSC, Systemic sclerosis; UC, ulcerative colitis; VIT: Vitiligo; MDD, major depressive disorder; SCZ, schizophrenia.

To evaluate the degree of heterogeneity necessary to achieve the observed genetic sharing for these autoimmune diseases, we calculated the GRS regression coefficient. We previously showed that the GRS regression coefficient approximates the expected heterogeneity proportion *π*^36^ assuming no pleiotropy. Based on the GRS coefficients, we observed *π* estimates ranging from 8-76% across the different autoimmune diseases in T1D and from 10-43% with RA (**Figure 4, Table 1**).

**Figure 4.**
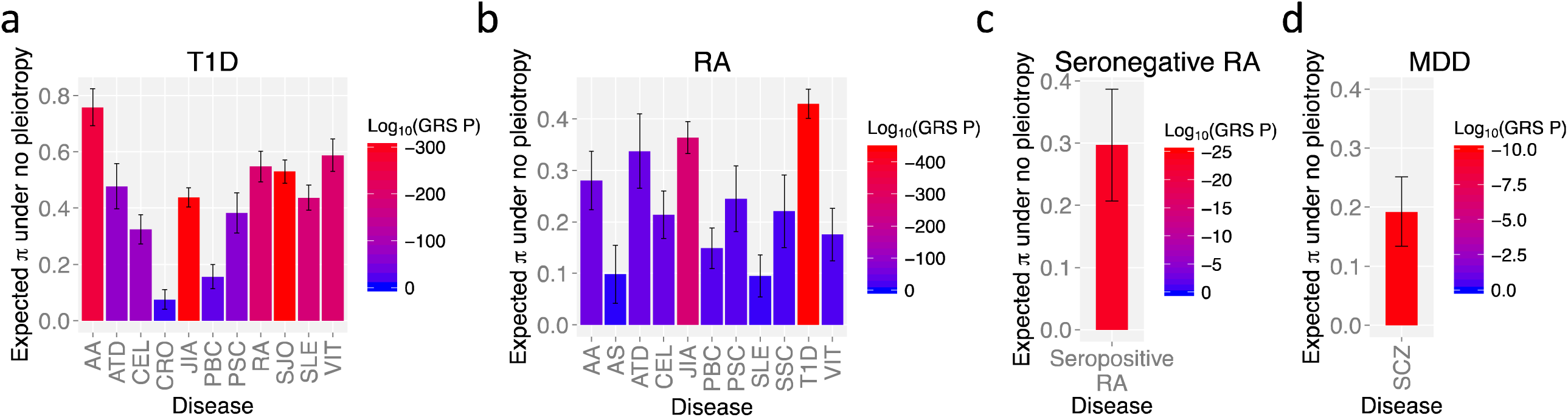
Genetic sharing between autoimmune diseases and psychiatric disorders. Out of 11 autoimmune diseases that have ≥10 pruned associated loci, only the diseases that have significant GRS P-values in positive directions are shown. Y-axis is the expected misclassifications if there is no pleiotropy, to explain observed genetic sharing. Vertical bars indicate 95% confidence intervals. Heterogeneity expected based on GRS analysis, assuming no pleiotropy for (a) T1D, (b) RA, (c) seronegative RA, and (d) MDD.

We then estimated the power of BUHMBOX to detect heterogeneity, using Bonferroni correction for 11 tests (p<4.5×10^−3^). We found that BUHMBOX is well powered for some autoimmune traits. Assuming *π*=0.2, four traits had >90% power for T1D, and four traits had >90% power for RA (**Figure 5**). Despite high power for certain traits, we observed no evidence of heterogeneity at all (corrected p>0.2; **Figure 6, Table 1**) suggesting that, for these autoimmune traits, genetic sharing is clearly due to pleiotropy and not heterogeneity. Autoimmune diseases share similar risk alleles and pathways with T1D and RA, and not by subgroups of genetically similar cases resulting from misclassifications or molecular subtypes.

**Figure 5.**
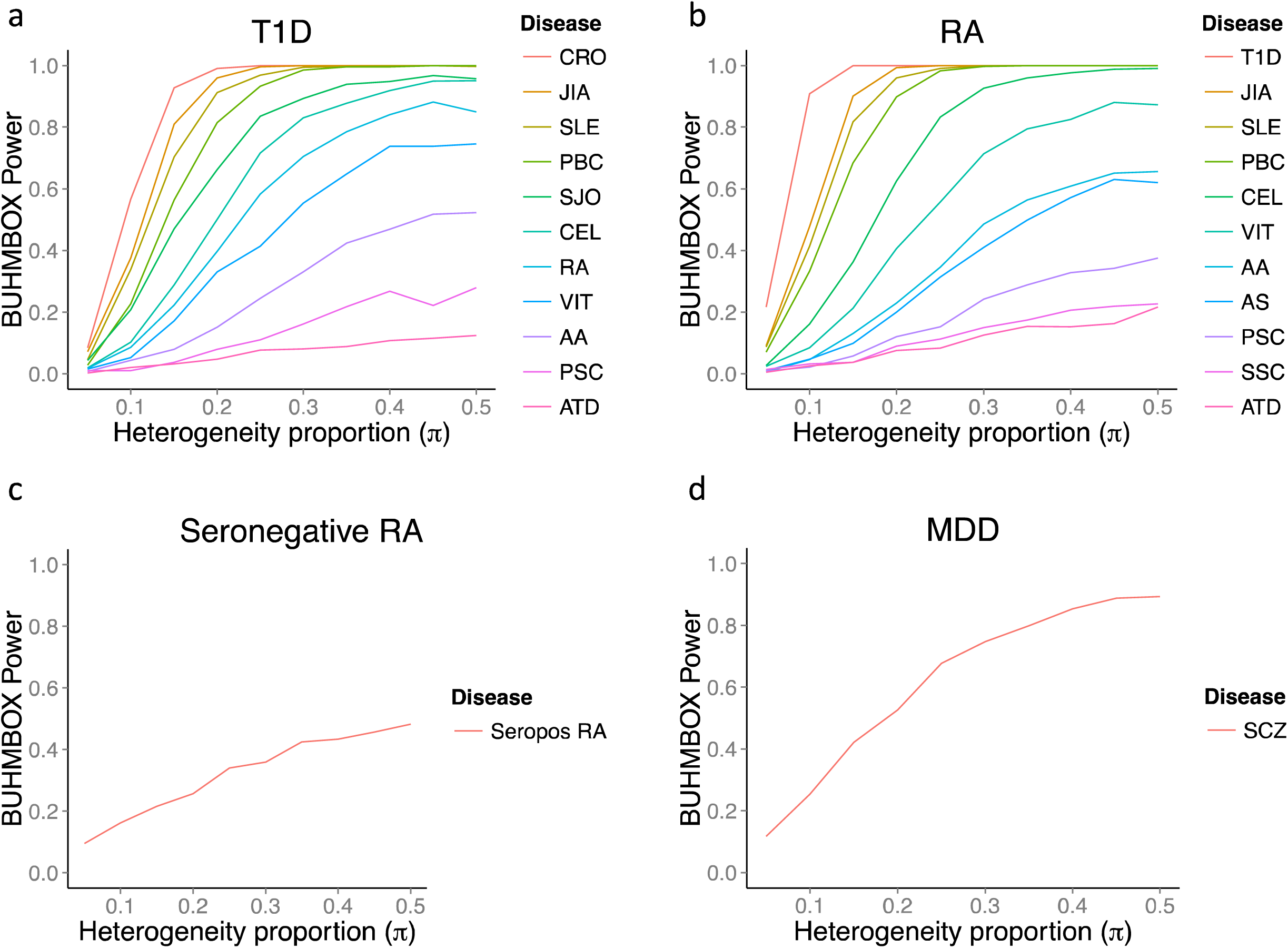
Statistical power of BUHMBOX to detect heterogeneity. Power was calculated by performing 1,000 simulations with corresponding sample size, number of risk alleles, risk allele frequencies, and adjusted odds ratios to account for pleiotropy. To calculate power for (c) and (d), we used a significance threshold of 0.05. For (a) and (b), the threshold was adjusted using the Bonferroni correction accounting 11 tests in T1D and RA, respectively.

**Figure 6.**
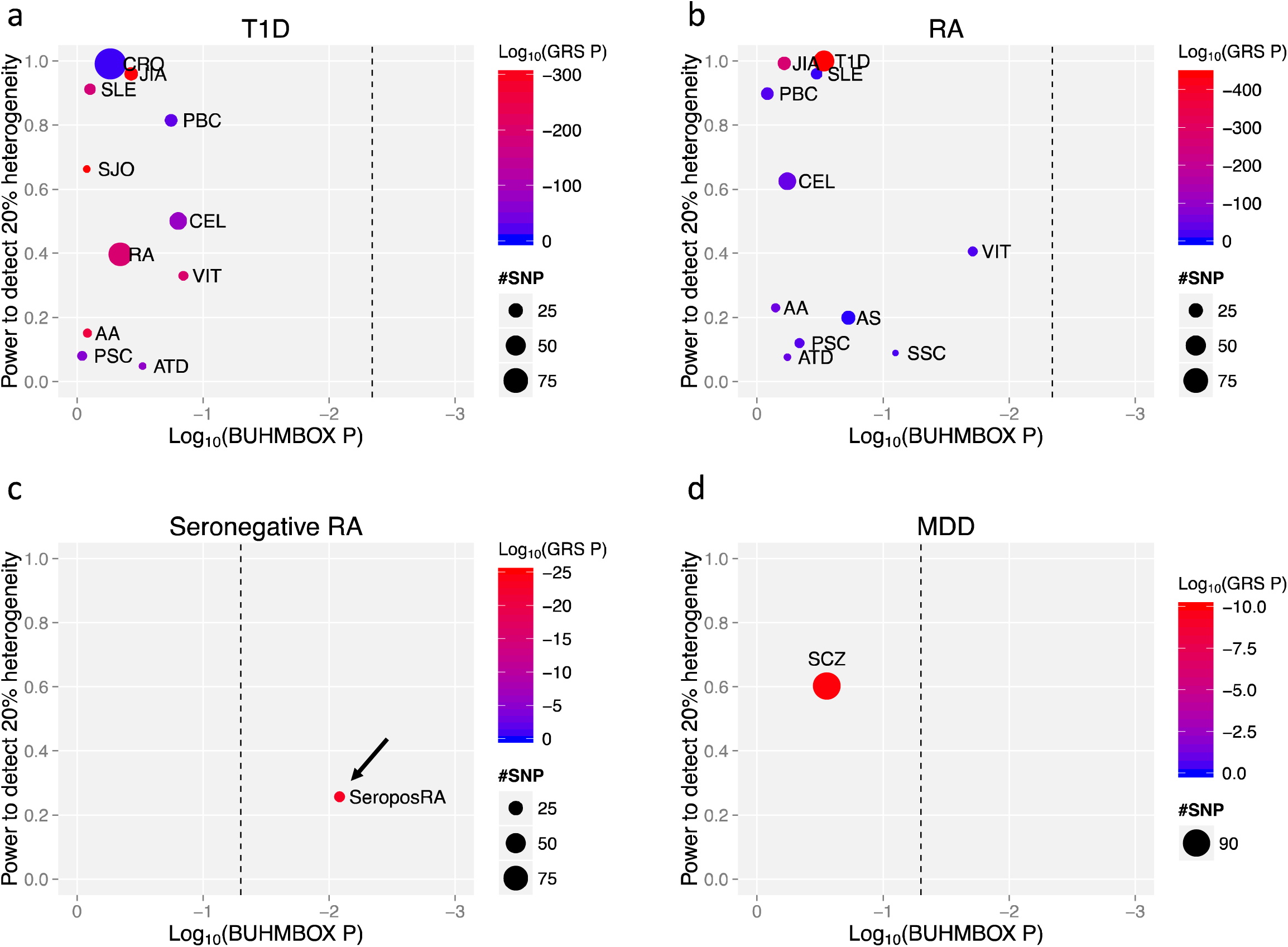
BUHMBOX results. Dashed vertical lines denote the Bonferroni-adjusted significance threshold.

### Application to subtype misclassifications in RA

RA consists of two subtypes, seropositive and seronegative, with distinct clinical outcomes and MHC associations^36^. These two subtypes are classified by whether patients are reactive to anti-CCP antibody. While anti-CCP testing is highly specific, it is not perfectly sensitive which results in some seropositive RA patients being misclassified as seronegative RA^18^. We previously demonstrated that there is shared genetic structure between seropositive and seronegative RA using the GRS approach^36^, which could imply misclassifications of up to 26.3% between the two RA subtypes.

We evaluated the ability of BUHMBOX to detect seropositive RA misclassifications in a seronegative RA cohort using only SNP data. We used the seronegative RA cohort (2,406 cases/15,870 controls) from the RA Immunochip Consortium^28^. Among 68 RA-associated independent loci, we chose SNPs that are associated to seropositive RA (p<5×10^−8^) but not seronegative RA (p>5×10^−8^) in our Immunochip data. This criterion resulted in 14 specific loci that are exclusively associated to seropositive RA (**Supplementary Table 3**). The seropositive RA GRS was significantly associated with seronegative RA case status (β=0.30, p=1.1×10^−23^). The regression coefficient (β=0.30) approximates the upper bound of the heterogeneity proportion *π* (**Figure 4**). Application of BUHMBOX suggested that coheritability was indeed explained by heterogeneity (p<0.008, **Figure 6, Supplementary Table 4**), consistent with potential subtype misclassifications.

### Application to major depressive disorder and schizophrenia

Current criteria for diagnosing psychiatric disorders reflect clinical syndromes, often with overlapping symptoms. As a result, psychiatric diagnoses for a patient may change as their symptoms evolves. MDD is thought to be a particularly heterogeneous psychiatric disorder, with relatively low diagnostic stability^19^. In addition to the potential for misdiagnosis, a subset of true MDD cases may be genetically more similar to schizophrenia. If heterogeneity with respect to schizophrenia risk alleles exists among MDD cases, then genetic studies would suggest evidence of coheritability between the two disorders^37^ as has been observed in previous studies^3, 6, 7^. The unintentional inclusion of “schizophrenia-like” MDD cases, due to diagnostic misclassification or genetically distinct subgroups, has been acknowledged and explored as a potential source of bias in coheritability studies by previous investigators^3, 37^.

We used BUHMBOX to test for a subgroup of “schizophrenia-like” cases in MDD. If a subset of MDD cases are misdiagnosed and in fact have schizophrenia, or are more genetically similar to schizophrenia, we would expect to see heterogeneity among MDD cases with respect to schizophrenia risk loci. We first evaluated evidence of shared genetic structure among 90 known schizophrenia associated loci^38^ (**Supplementary Table 3**) in 9,238 MDD cases and 7,521 controls from the Major Depressive Disorder Working Group of the Psychiatric Genomics Consortium^39^ (see **Supplementary Table 5** for details of the MDD dataset). Consistent with previous findings^3, 6^, the GRS based on these loci was significantly associated with MDD case status (p=1.54 × 10^−5^) indicating shared genetic structure between schizophrenia and MDD (**Figure 4**). For the GRS analysis we used a refined subset of the total sample (6,382 MDD cases and 5,614 controls), which excluded samples that overlapped with the schizophrenia GWAS^38^ (**Methods**). The BUHMBOX p-value was not significant (p=0.28), indicating no excess positive correlations among schizophrenia loci within MDD cases (**Figure 6, Supplementary Table 4**). Our findings suggest no evidence of a subgroup of schizophrenia-like MDD cases. However, we note that we did not have adequate statistical power to detect heterogeneity in the context of a small degree of heterogeneity. Given the MDD sample size and the number of currently known schizophrenia risk loci, there was 53% power to detect 20% heterogeneity, but only 25% power to detect 10% heterogeneity (**Figure 5**).

## DISCUSSION

Here we present BUHMBOX, which can distinguish whether shared genetic structure between two traits is the consequence of heterogeneity versus pleiotropy based on SNP genotype data alone. Our method builds upon recent observations emerging in the literature of shared genetic structures in autoimmune, neuropsychiatric, and metabolic diseases. BUHMBOX utilizes the intuition that if heterogeneity exists, independent loci will show non-random positive correlations; importantly, we correct for population structure and long-range LD, which may serve as confounders for this analysis. Heterogeneity can be caused by (1) misdiagnosis, (2) a subgroup of cases that share molecular etiology with another disease, or (3) an excessive number of cases affected by comorbidity compared to what would be expected under pleiotropy alone, which can happen because of ascertainment bias or causal relationships between diseases (i.e. mediated pleiotropy in a subgroup of cases). We emphasize that it is critical to appropriately interpret the source of heterogeneity, which will depend on the biological and clinical relationship between the two traits. We provide detailed information to guide interpretation in the **Supplementary Information.**

We demonstrated that much of the genetic sharing between autoimmune diseases is due to pleiotropy. We do note that for a few traits we had modest power (**Figure 5**) to detect heterogeneity proportions less than *π*=0.2. One exception was our analysis that suggested that seronegative RA samples might contain misclassified seropositive RA cases. In contrast we were underpowered to draw a definitive conclusion as to whether a subset of MDD cases are genetically similar to schizophrenia cases, although undoubtedly as MDD cohorts increase in size we will be able to reassess more accurately whether smaller proportions of heterogeneity might partially explain the observed coheritability. Our current results are in line with the results of an analytical study^37^, which concluded that the observed degree of pleiotropy between psychiatric diseases is unlikely explained by misclassifications alone.

We have shown that the power of BUHMBOX is a function of sample size, number of loci, effect sizes and allele frequencies of loci, and the heterogeneity proportion *π.* For detecting subtle heterogeneity (*π* <0.1), current datasets were often not well powered. But, we expect that in future studies, as we increase the sample size as well as the number of known associated loci, our method will become increasingly powerful for detecting subtle heterogeneity. Even with existing genetic data, a potential strategy to augment power is to include a larger number of SNPs selected using less stringent significance thresholds, an approach referred to as polygenic modeling^3, 10, 11^. We performed simulations to demonstrate that polygenic modeling can indeed increase the power substantially (**Supplementary Methods** and **Supplementary Figure 2**).

We designed BUHMBOX to identify the presence of heterogeneity, in the situation where we do not know the specific membership of individuals to the subgroup. In this paper, it was not our goal to uncover subgroup membership using genetic data, because genetic information is typically not adequate to clearly classify individuals into subgroups. In certain situations, we may be able to discern membership. For example, for the misclassification of seropositive RA samples in the seronegative RA cohort, as serological assays advance we will have a means to more precisely define membership^40^. If we know the membership, it is possible to perform additional analyses such as comparing GRS between subgroups.

When comparing BUHMBOX to existing approaches, we focused on the GRS method. However, the results of our comparison also apply to other existing methods such as mixed-model-based approaches^5, 6^ and LD-score-based approaches^7^, which are similar to the GRS approach in the sense that they detect both pleiotropy and heterogeneity. We expect that BUHMBOX will complement any of these methods to facilitate interpretation of observed genetic sharing between traits. More broadly, BUHMBOX can be thought of as capturing a specific form of epistasis where risk alleles correlate positively within the additive model. Our statistical approach may therefore be extended to have application beyond heterogeneity, including identification of missing heritability resulting from clinical heterogeneity^41^. These applications will become more feasible as functional annotations of SNPs advance in the coming years.

## ACKNOWLEDGMENTS

This work is supported in part by funding from the National Institutes of Health (1R01AR063759 (SR), 5U01GM092691-05 (SR), 1UH2AR067677-01 (SR), U19 AI111224-01 (SR)) and the Doris Duke Charitable Foundation Grant #2013097. JGP is supported by Fulbright Canada, the Weston Foundation, and by Brain Canada through the Canada Brain Research Fund. KS is supported by an NIH training grant (T32 HG002295). This research utilizes resources provided by the Type 1 Diabetes Genetics Consortium, a collaborative clinical study sponsored by the National Institute of Diabetes and Digestive and Kidney Diseases (NIDDK), National Institute of Allergy and Infectious Diseases (NIAID), National Human Genome Research Institute (NHGRI), National Institute of Child Health and Human Development (NICHD), and Juvenile Diabetes Research Foundation International (JDRF) and supported by U01 DK062418.

## AUTHOR CONTRIBUTIONS

BH and SR conceived the statistical approach and organized the project. BH, JGP and SR led and coordinated analyses and wrote the initial manuscript. ES and NW provided guidance on the statistical approach. KS, CHL, DD, XH, YRP, and EK contributed to the implementation of specific analyses and offered feedback to the statistical methodologies. PKG, SRD, JW, JM, SE, LK, SR and TH contributed RA samples and insight on the clinical implications to RA. W-M C, S O-G, and SSR contributed T1D samples and insight on clinical implications to T1D. MDDWG contributed MDD samples and insight on the clinical implications to MDD. All authors contributed to the final manuscript.

## ONLINE METHODS

### Genetic risk score approach

Given *M* independent risk loci associated to D_B_, we calculated the GRS of individual *i* as

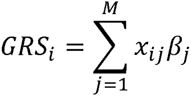

where *x_ij_* is individual *i*’s risk allele dosage at marker *j,* and *β_j_* is the effect size (log odds ratio) of risk allele at marker *j* for disease D_B_. The GRS approach calculates GRSs for all individuals and associates GRSs to the case/control status of D_A_.

### The BUHMBOX approach

To detect heterogeneity, we developed the following procedure:

1. Prune SNPs associated with D_B_ based on control LD (excluding SNPs that are r^2^>0.1 or within ±1Mb to other SNPs)
2. Obtain risk allele dosages of pruned SNPs from D_A_ cases and controls
3. Regress out PCs from risk allele dosages to obtain residual dosages
4. Calculate R, correlation matrix of residual dosages in *N* cases with D_A_ and R’, using *N’* of controls
5. Calculate delta-correlations:

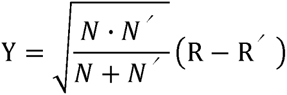
6. Calculate the BUHMBOX statistic:

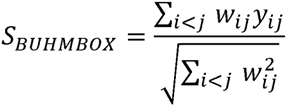

where *y_ij_* is the element in *Y* at row *i* and column *j.* Given *M* pruned SNPs, (*i,j*) iterates *M(M-1)/2* non-diagonal elements of *Y*. The *w_ij_* term is a weighting function that is designed to maximize power, such that (equation (13) in Supplementary Methods):

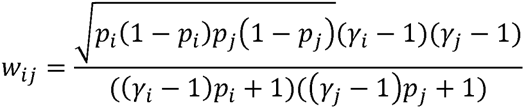

where *p_i_* is RAF of SNP *i,* and *γ_i_* is the OR of SNP *i* for D_B_. The BUHMBOX statistic follows *N*(0,1) under the null hypothesis, and is one-sided in the positive direction. Thus, the p-value is *p_BUHMBOX_* = 1 ^_^ Φ (*S_BUHMBOX_*) where *Φ* is the cumulative density function of the standard normal distribution. In the context of heterogeneity, excessive positive correlations among D_B_ risk alleles in D_A_ cases result in *p_BUHMBOX_* < *α*. See **Supplementary Table 1** for comparison of BUHMBOX and GRS approaches. The BUHMBOX test statistic was inspired by previous work deriving covariance between correlation estimates^42^ and on combining dependent estimates.^43^ For details of the intuition, derivation, optimization, and interpretation of the BUHMBOX test statistic, see **Supplementary Information**.

### Code availability

BUHMBOX has been fully implemented as a publicly available R script (https://www.broadinstitute.org/mpg/buhmbox/).

### Power and false positive rate simulations

Given sample size of D_A_ cases (*N*), number of risk loci associated to D_B_ (*M*), proportion of D_A_ cases that actually show genetic characteristics of D_B_ (heterogeneity proportion *π*), we simulated studies to estimate power of our method as follows. To simulate a reasonable joint distribution of RAFs and ORs, we downloaded the GWAS catalog (as of 29 April 2014). Among all binary traits in the catalog, we selected traits with ≥50 reported SNPs resulting in 22 traits with 1,480 SNPs. From these SNPs, we sampled *M* pairs of RAF (*p*) and their corresponding OR (*γ*). To simulate genotypes, we set the RAF of a subgroup (*Nπ* individuals) to *γp/((γ-1)p+1)* and *p* for the other subgroup *(N(1-π)* individuals), because *Nπ* individuals can be thought of as D_B_ cases. Within each subgroup, we generated genotypes assuming that the risk alleles are distributed according to the Hardy-Weinberg equilibrium (HWE) and that the risk loci are independent. We assume HWE in cases because we assume an additive disease model. Then we applied our method to calculate the p-value. We repeated this 1,000 times to approximate power as the proportion of the repeats whose p-values were ≤0.05. We evaluated power for different values of *N*, *M*, and *π.*

Under the assumption that the loci are independent, the false positive rate simulation was equivalent to the power simulation described above with the only difference being that *π* was set to zero, which forced the null hypothesis. We measured false positive rate by assuming *N*=1,000 and *M*=20, and constructing 1,000,000 such studies.

### Linkage disequilibrium simulations

To simulate realistic LD, we used chromosome 22 data from control individuals in the Swedish EIRA cohort of the RA dataset (2,762 cases/1,940 controls)^28^. Then, we assigned half of control individuals as cases and the rest as controls. To generate 1,000 random datasets, we thinned the data by 10-fold with different seed numbers using PLINK^44^ (with the command --thin 0.1). We then pruned each of the 1,000 datasets using PLINK^44^ with r^2^ criterion of 0.5 or 0.1.

### Population stratification simulations

To simulate population stratification, we combined HapMap^29^ release 23 data (60 CEU founders, 60 YRI founders, and 90 JPT+CHB founders). We set CEU+YRI as cases and JPT+CHB as controls. We calculated PCs after LD pruning r^2^<0.1. We randomly selected 5,000 sets of 22 independent SNPs, each of which was selected from each autosome. We also combined a Northern Europe RA cohort (Swedish EIRA; 2,762 cases/1,940 controls) and a Southern Europe cohort (Spain; 807 cases/399 controls) from the RA dataset^28^. Similar to the linkage disequilibrium simulation, we thinned the chromosome 22 data by 10-fold using 1,000 different random seeds, and applied pruning with criterion r^2^<0.1.

### Application to specific phenotypes

*Type 1 diabetes dataset.* To evaluate pleiotropy and heterogeneity between 18 autoimmune diseases and T1D, we applied GRS and BUHMBOX approaches to the UK case-control dataset provided by the Type 1 Diabetes Genetics Consortium^35^, which consisted of a total of 16,086 samples (6,670 cases and 9,416 controls) from three collections: (1) cases from the UK-GRID, (2) shared controls from the British 1958 Birth Cohort and (3) shared controls from Blood Services controls (data release February 4, 2012, hg18). The samples were collected from 13 regions. All samples were collected after obtaining informed consent, and were genotyped on the ImmunoChip array. GRS and BUHMBOX analyses were conducted using the region index as covariates.

*Rheumatoid arthritis dataset.* To evaluate pleiotropy and heterogeneity between 18 autoimmune diseases and RA, we used the RA Immunochip consortium data from six RA case-control cohorts (UK, US, Dutch, Spanish, Swedish Umea, and Swedish EIRA)^28^. To evaluate pleiotropy to autoimmune diseases, we used 7,279 seropositive RA cases and 15,870 controls. To evaluate misclassifications of RA subtypes, we used 2,406 seronegative RA samples and the same controls. Seropositive and seronegative RA patients were defined in each cohort using standard clinical practices to assess whether patients were reactive to anti-CCP antibody^36^. All samples provided informed consent, and were collected through institutional review board approved protocols. All individuals self-reported as white and of European descent. Samples were genotyped with the Immunochip custom array. We merged the data of six cohorts into one, adding binary variables indicating cohorts as covariates in the analysis.

*Defining autoimmune risk loci.* Immunobase curations (http://www.immunobase.org/downloads/regions-files-archives/2015-06-07/*assoc_variantsTAB; accessed 7 June 2015) available for 18 autoimmune diseases were used to define genome-wide significant risk loci. We did not include inflammatory bowel disease, due to its redundancy with Crohn’s disease and ulcerative colitis. For each of the 18 autoimmune diseases analyzed we pruned the list of index SNPs obtained from Immunobase in PLINK^44^ with options --r2 --ld-window-r2 0.1, using the 1000 Genomes Phase 1 European reference panel for LD. For all pairs of SNPs with r^2^ > 0.1, we kept the most strongly associated SNP. To ensure completely independent risk loci we also removed SNPs annotated as being located in the same chromosomal region in Immunobase, again keeping the most strongly associated index SNP (**Supplementary Table 3**). When a locus was not in the Immunochip datasets, we looked for proxy (r^2^>0.2) based on the 10000 Genomes data.

*Major depressive disorder dataset.* We used BUHMBOX to investigate the relationship between MDD and schizophrenia, which have been previously reported to share genetic etiology based on polygenic risk scoring^3^ and coheritability analyses^6^. The full MDD sample analyzed comprised nine GWAS datasets collected from eight separate studies (**Supplementary Table 5**) as previously described^39^. All samples were collected through institutional review board approved protocols were collected with consent. For the GRS analysis, individual MDD samples (four cases, 886 controls) that overlapped with those in the schizophrenia GWAS^38^ were removed from the analysis; three GWAS cohorts with an insufficient number of independent control samples (n < 5) were also removed from the analysis. GRS analyses were conducted in each of the remaining six GWAS datasets (**Supplementary Table 5**), followed by meta-analysis of the GRS. To obtain the overall ß GRS effect size and test statistic we used the inverse-variance weighted fixed effects method. For BUHMBOX, we used the full dataset; analyses were conducted in each of the nine GWAS datasets (**Supplementary Table 5**) followed by meta-analysis. Because the BUHMBOX statistic is a z-score, we meta-analyzed BUHMBOX results across the datasets using the standard weighted sum of z-score approach, where z-scores are weighted by the square root of the sample size.

*Defining schizophrenia risk loci.* Schizophrenia associated SNPs were selected as those showing genome-wide significant association with schizophrenia (p < 5×10^−8^) in the recent GWAS mega-analysis by the Psychiatric Genomics Consortium^38^. For schizophrenia associated SNPs not directly genotyped in the MDD GWAS datasets, we selected proxy SNPs as those with the highest r^2^ from the list of all proxies with r^2^ > 0.20 using the 1000 Genomes Phase 1 European reference panel. Of the 97 schizophrenia associated SNPs (11 indels were not considered in our analysis), 90 LD-independent SNPs (r^2^ > 0.1, distance > 1Mb) were available for analysis in the MDD GWAS datasets either via direct genotyping or a proxy (see **Supplementary Table 3** for a detailed list of SNPs).

